# Reciprocal recombination reflects sexual reproduction in symbiotic arbuscular mycorrhizal fungi

**DOI:** 10.1101/2021.03.05.434083

**Authors:** Ivan D. Mateus, Ben Auxier, Mam M. S. Ndiaye, Joaquim Cruz, Soon-Jae Lee, Ian R. Sanders

## Abstract

Arbuscular mycorrhizal fungi (AMF) are part of the most widespread fungal-plant symbiosis. They colonize at least 80% of plant species, promote plant growth and plant diversity. These fungi are multinucleated and contain either one or two haploid nuclear genotypes (monokaryon and dikaryon) identified by the alleles at a putative mating-type locus. This taxon has been considered as an ancient asexual scandal because of the lack of observable sexual structures. Despite identification of a putative mating-type locus and functional activation of genes related to mating when two isolates co-exist, it remains unknown if AMF take part in a mainly sexual life cycle.

We used publicly available genome sequences to test if dikaryon genomes display signatures of sexual reproduction in the form of reciprocal recombination patterns, or if they display signatures of parasexual reproduction involving gene conversion.

We used short-read and long-read sequence data to identify nucleus genotype-specific haplotypes within dikaryons and then compared them to orthologous gene sequences from related monokaryon isolates displaying the same putative MAT-types. We observed that these genotype-specific haplotypes display reciprocal recombination and not gene conversion.

These results are consistent with a sexual origin of the dikaryon rather than a parasexual origin and provides an important step to understand the life cycle of these globally important symbiotic fungi.

## Introduction

Arbuscular mycorrhizal fungi are plant symbionts, forming symbioses with most plant species, promoting plant growth (Harrison, 1997), plant community diversity (van der Heijden *et al.*, 1998; Antunes *et al.*, 2011) and how plants cope with biotic (Thygesen *et al.*, 2004) or abiotic stresses (Augé, 2001). As a consequence, they are widely used in agriculture (Smith & Read, 2008). In a single agricultural field, the presence of at least 17 different genotypes of *Rhizophagus irregularis* displaying spatial genetic structure have been detected (Koch *et al.*, 2004; Croll *et al.*, 2008). Furthermore, these *R. irregularis* isolates display different levels of within isolate genetic diversity (Wyss *et al.*, 2016), which has been reported to produce differential effects on plant growth (Angelard *et al.*, 2010). Understanding how genetic variability is generated in AMF, is important because it could be harnessed to generate genetic variants that could be beneficial for their plant hosts (Sanders, 2010).

AMF are part of the Glomeromycotina subphylum (Spatafora *et al.*, 2016), which fossil records date to at least ~400 Million years ago (Remy *et al.*, 1994). They are coenocytic (without septa separating otherwise adjacent compartments), their hyphae harbor hundreds of nuclei within the same cytoplasm (Marleau *et al.*, 2011) and no single-nucleus state has been recorded. The nuclei of these fungi have been reported as haploid (Lin *et al.*, 2014; Ropars *et al.*, 2016; Kobayashi *et al.*, 2018; Morin *et al.*, 2019). This group of fungi has been previously considered as an ancient asexual scandal (Judson & Normark, 1996), due to low morphological diversification and the absence of observable sexual structures. However, evidence suggests that sexual reproduction could be possible in AMF since these fungi contain a complete meiosis machinery (Halary *et al.*, 2011). Furthermore, a putative mating-type determining locus (MAT) has been proposed (Ropars *et al.*, 2016), population genetic data suggests the existence of recombination in AMF populations (Croll & Sanders, 2009) and activation of genes related to meiosis has been detected when different isolates of the same species co-exist in plant roots (Mateus *et al.*, 2020, 2021).

Several *R. irregularis* isolates issued from the same geographic location have been reported as monokaryon (A1,B12,C2) and dikaryon (A4, A5, C3 and G1) (Ropars *et al.*, 2016; Chen *et al.*, 2018a; Masclaux *et al.*, 2018; Kokkoris *et al.*, 2021), evidencing that monokaryon and dikaryon isolates co-exist in the same location. Single-nucleus from dikaryons (isolates A4, A5 and SL1) cluster into two genetically different nucleotypes (where nucleotype refers to the genotype of a nucleus), that are associated with the identity of a putative MAT-locus (Chen *et al.*, 2018a). This demonstrates that the presence of two copies of the putative MAT-locus is a reliable marker of the dikaryon state.

Like most fungi, AMF can undergo anastomosis, the fusion of hyphae. Through these connections, bi-directional flow of cytoplasm has been observed between genetically different AMF individuals (Giovannetti *et al.*, 1999). Via anastomosis, the transfer of genetic material between vegetative compatible isolates (parasexuality) has been suggested as a mechanism of maintenance of genetic diversity in the absence of sexual recombination (Bever & Morton, 1999). In fungi, hyphal fusion between different individuals leads to cell death, however, non-self vegetative compatibility has also been observed in AMF when different isolates form perfect hyphal fusions (Croll *et al.*, 2009). In the model species *Aspergillus nidulans* parasexuality involves fusion of two haploid nuclei, mitotic recombination and haploidization of the diploid nuclei by i.e. chromosome loss (Pontecorvo, 1956). In consequence, a genomic signature of parasexuality is gene-conversion where there is a loss of heterozygosity. However, sexual reproduction can also display signatures of gene-conversion but also involves meiotic recombination and the segregation on reciprocal products.

The existence and relevance of sexuality and/or parasexuality for the evolution of AMF remains unknown (Yildirir *et al.*, 2020). It is still unknown whether the transition between a dikaryon and monokaryon life stages involves a sexual event involving meiotic recombination, or a parasexual event.

In the absence of stable transformation methods in AMF (Helber & Requena, 2008), haplotype analyses of genomic data could be an important resource to identify genomic signatures of mitotic or meiotic recombination in dikaryon isolates. The haplotypes from dikaryon isolates can then be compared to orthologous sequences of related monokaryon isolates (that share the same MAT-type) and could allow to identify genomic signatures of a sexual event, involving meiotic recombination, or a parasexual event which results in the loss of heterozygosity or gene conversion.

AMF genomes display large gene duplication events (Morin *et al.*, 2019), which make it difficult to call haplotypes. The identification of haplotypes in dikaryon isolates could be made by: 1) analyzing short-read sequences to identify genome-wide copy number variation. These analyses consist in obtaining the read depth, or coverage, after mapping the reads to a genome assembly and identifying changes in coverage across the genome (Yoon *et al.*, 2009). In AMF, a drop in coverage analysis was used to originally identify the putative MAT-locus (Ropars *et al.*, 2016) highlighting the potential to identify nucleotype-specific haplotypes in dikaryons with this technique. 2) Analyzing long-read sequencing to identify haplotypes and, consequently, nucleotype-specific haplotypes in fungal dikaryon isolates (Li *et al.*, 2019).

Here, we demonstrated that AMF dikaryons display reciprocal recombination by analyzing nucleotype-specific haplotypes in dikaryon (A5) and monokaryon isolates (A1-C2). In this study we used publicly available whole genome, single-nucleus short-read sequence data and long-read bulk-isolate sequence data to identify nucleotype-specific haplotypes in dikaryon isolates. We identified regions displaying drops in coverage in short-read whole genome sequence assemblies. In these regions we detected the presence of genes that have two copies in the dikaryon isolates and one copy in monokaryons. We then confirmed independently, with short-read genome sequence data from single-nucleus, that in dikaryon isolates, different nuclei have different alleles and that they are not always associated to the MAT identity, evidencing a reciprocal recombination genomic signature. Finally, we validated the analysis, by evaluating long-read genome sequence assemblies and confirmed that A5 dikaryon display genomic signatures of reciprocal recombination, which suggest sexual reproduction in AMF.

## Materials and Methods

### Source data

We used public-available sequence reads, genome assemblies and annotations of isolates A1, A4, A5, C2 of *R. irregularis* for this study, including data from bulk-isolate and single-nucleus sequencing (Supplementary Table 1). We used short-read whole genome assemblies from isolates A1, A4, A5 and C2 (Ropars *et al.*, 2016). We used single-nucleus genome assemblies from isolates A1, A5 and C2 (Chen *et al.*, 2018a). We also analyzed long-read genome assemblies of isolates A1, A5 and C2 (Chaturvedi). We downloaded the sequence reads from the sequence read archive (SRA) using the SRAtoolkit software with the prefetch and fastq-dump tools (Leinonen *et al.*, 2011).

### Coverage analysis

We first trimmed the sequence reads using Trim Galore! (Krueger, 2015) with the default parameters. We then used BWA (Li, 2013) to index the reference genome assemblies and BWA mem -M (Li, 2013) to map the reads to the reference whole-genome assemblies. We mapped the reads coming from a given isolate to the reference genome assembly of the same isolate (i.e. reads A1 mapped to reference A1). We then kept the reads that display a mapping quality of at least 30. We used the genomecov tool from bedtools (Quinlan & Hall, 2010) to calculate the coverage for each position. We then created a ready-to-use algorithm that detects genome-wide drop in coverage analysis in whole-genome data (Supplementary File 1). The algorithm divides the data in portions of 50kb. Then, with a sliding window approach consisting of windows of 400bp and steps of 100bp, the algorithm searches for drops in coverage of 0.3-0.6 times lower than the median coverage of the entire genome and that with a minimum length of 1000bp (Please refer to Supplementary File 1 for the algorithm specifications implementated in the R programming language). We then further filtered these drop in coverage regions by keeping only the regions that display an average of 1.25 coverage difference between the neighboring regions and the drop in coverage region.

### Gene detection in drops

We identified all the genes located within genomic regions that presented a drop of sequencing coverage. We used the ‘intersect’ command from the BEDTools suite with the existing gene annotations corresponding to each *R. irregularis* isolate (GTF format) and their query regions with drops in coverage (BED format) to identify overlapping genes (Quinlan & Hall, 2010). Genes in scaffolds smaller that 1kb were not considered for further analyses.

### *de novo* single-nucleus assemblies

We trimmed the raw reads by using TrimGalore-0.6.0 (Krueger, 2015) with default parameters. After trimming, we performed single-nucleus *de novo* assemblies with SPAdes v3.14 (Bankevich *et al.*, 2012) with the following parameters: -k 21,33,55,77 --sc --careful --cov-cutoff auto. The resulted single-nucleus genome assemblies were used for further analysis. The length, number of contigs and N50 value of the *de novo* assemblies was evaluated with quast-5.1.0rc1 with default parameters (Mikheenko *et al.*, 2018).

### Identification of genes in genome assemblies

To identify the position of gene sequences on the different genome assemblies, we first extracted a query sequence. We then used the console NCBI+ blast suite (Camacho *et al.*, 2009) to blast the query against the desired target. In the case of the putative MAT-locus, we used the homeodomain genes HD2 and HD1-like as query (HD2:KT946661.1, HD1-like: KU597387 from isolate A1). For further downstream analyses, we extracted the sequences from the genome assemblies by using the blastdbcmd command from the NCBI+ suite. We used a reciprocal blast approach to identify the gene sequences corresponding between the whole genome sequence data and the single-nucleus data. We considered the bests hits by evaluating the % identity, mismatches, e-value and bitscore.

### Orthology inference

We used Orthofinder 2.3.11 (Emms & Kelly, 2019) to identify orthologs of genes found inside the drop in coverage regions within the same isolate. We also identified orthologs in the long-read assemblies and identified single copy orthologs in isolates A1, C2, the primary assembly of A5 and the haplotig assembly of A5. We used the orthogroups output from Orthofinder for the different analyses.

### Synteny plots

We compared genomic regions by performing synteny plots computed with EasyFig2.2.3 (Sullivan *et al.*, 2011). We provide full Genbank files to compare genomic regions to each other. The software executes a blast comparison between the regions to determine their homology.

### Genetic distance between nucleus specific haplotypes

Coding sequences for the 12 confirmed nucleotype-specific genes were extracted using the Blast+ command line blastdbcmd tool (Camacho *et al.*, 2009). The sequences were then aligned with MAFFT (Katoh *et al.*, 2017) using the --auto option. Then, the ape package (Paradis *et al.*, 2004) of R was used to calculate the pairwise distance between the 4 alleles (2 from A5, and 1 each from A1 and C2).

### Recombination detection

We compared the sequences from drop in coverage regions from both nuclei specific haplotypes of isolate A5 and isolates A1 and C2 to detect if isolate A5 display recombination events between the two putative parental isolates. After identification of the syntenic region among the different isolates, we aligned the sequences with MAFFT (Katoh *et al.*, 2017) and evaluated whether the sequence of one of the nucleotype of isolate A5 was similar to A1 and the other similar to C2.

### Phylogenetic analyses

We used MEGA-X (Kumar *et al.*, 2018) for the different phylogenetic reconstructions shown in the study. We first aligned the data with ClustalW. We then find the best DNA models describing the relation between the sequences. Finally, we used a maximum likelihood phylogeny reconstruction with 100 bootstraps to infer the phylogenetic relation among the samples. In several cases, we were not able to perform maximum likelihood phylogenies because of the low number of samples to compare, so UPGMA trees were done instead. Phylogenetic reconstructions of the different orthologous groups on Figure 4 where produced by the Orthofinder software.

## Results

### Drop in coverage analysis reveals potential nucleotype-specific haplotypes

Previously, a drop in coverage analysis of selected regions was used for the identification of a putative MAT-locus in *R. irregularis* (Ropars *et al.*, 2016). To perform a similar analysis across the entire genome, we developed a script (Supplementary File 1) that allows us to identify genome-wide drop in coverage events (for the accessions of raw data and genome assemblies used in this study see Supplementary Table 1).

We identified drops in coverage in 4 different isolates of *R. irregularis* which are reported to be dikaryons (A4 and A5) and monokaryons (A1 and C2) (Figure 1, Supplementary Table 2). The number of coverage drops and genes inside the drop in coverage regions were different between dikaryons (A4-A5) and monokaryons (A1-C2) (Supplementary Table 3). These results indicated that dikaryotic isolates displayed more heterozygous regions than the monokaryons, suggesting that the genes present within the regions showing a drop in coverage are potential candidates for highly divergent, nucleotype-specific alleles in dikaryons. Confirming the reliability of our approach, we detected the expected drop in coverage in the putative MAT-locus region in isolates A4 and A5 but not in isolates A1 and C2 (Figure 1d).

**Figure 1.**
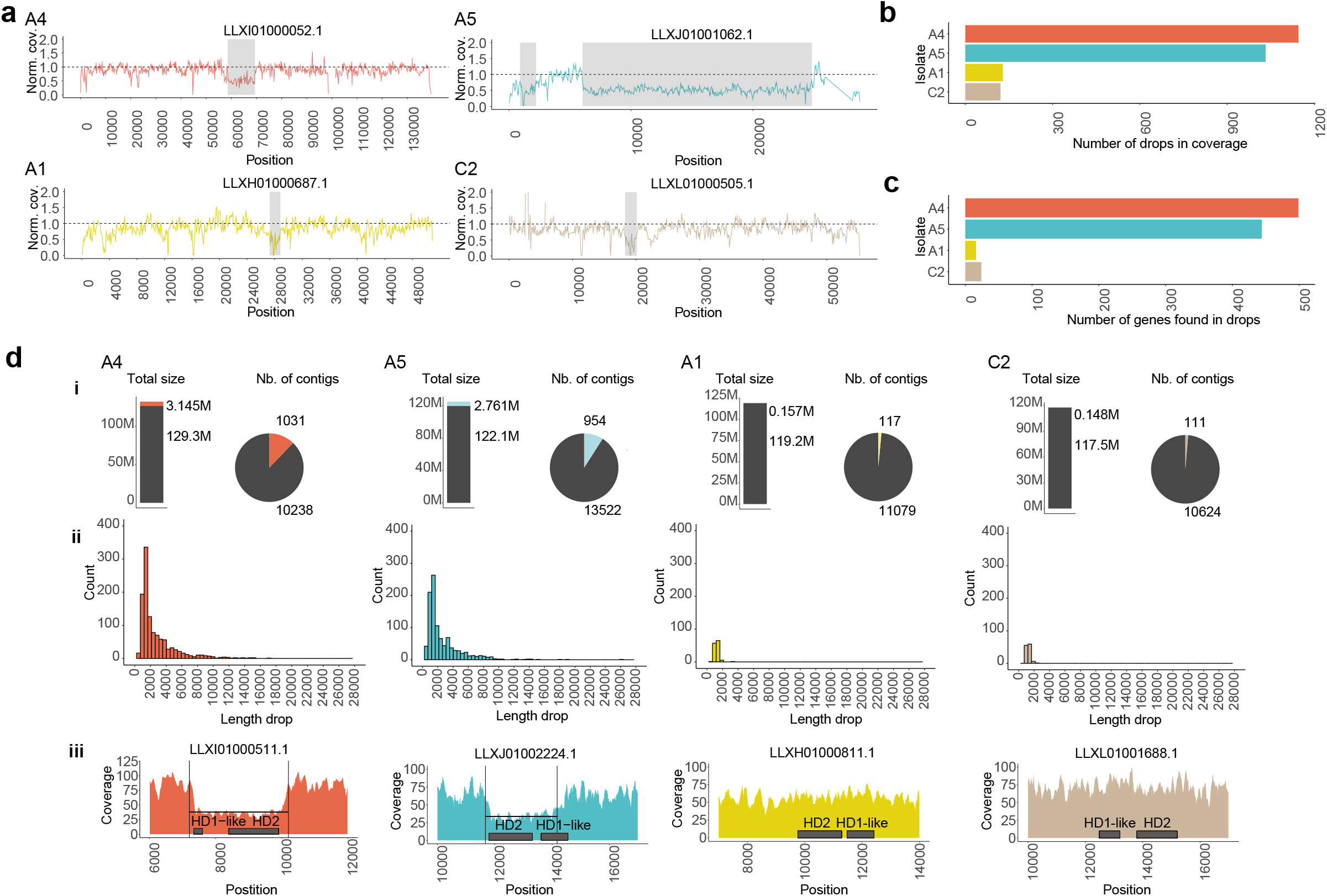
Drop in coverage events in isolates A4, A5, A1 and C2. **a,** Examples of drop in coverage events. We plotted the normalized coverage (y) per position (x). Grey rectangles represent the region detected by the algorithm. The horizontal dashed line represents the normalized coverage. **b,** Number of regions showing a drop in coverage that were detected in each isolate. **c,** Number of genes found in the regions showing a drop in coverage in each isolate. **d,** Summary statistics of regions showing a drop in coverage: i, proportion of total length of regions showing a drop in coverage and proportion of contigs that contain regions with a drop in coverage. ii, Histogram representing the lengths of identified regions where a drop in coverage was detected. iii, Coverage plot on the putative MAT-locus. Drop in coverage was detected in isolates A4 and A5 but not in A1 and C2.

One cause for a drop in coverage could be copy number variation between the nucleotypes in a dikaryon. To test for this, we inferred orthologous gene families among the different isolates to identify if the genes present in the drop in coverage regions displayed more than one copy in their own genome. We used the gene annotation available for each isolate and inferred the orthology of all the genes present in each genome. Many orthologous groups had more than one copy within each isolate (A4: 20%, A5: 18%, A1: 17% and C2: 19%; Supplementary figure 1, Supplementary Table 4) consistent with the high reported incidence of paralogs in these fungi (Morin *et al.*, 2019). We further identified orthologous gene families of genes detected in drop in coverage in isolates A4 and A5 independently. Under the assumption of a monokaryon-dikaryon genome organization in *R. irregularis*, to avoid the confounding effect of duplications and reduce the complexity of the dataset, we retained only the orthologous groups that are present in the drop in coverage regions and that display two copies in the dikaryon isolates (A4, A5) and a single copy in the monokaryons (A1, C2) (Figure 2a, Supplementary Table 5). In the regions where a drop in coverage was detected, we identified 32 orthologous groups that are present with two copies in isolate A4 and only a single copy in isolates A1 and C2. We also identified 27 orthologous groups in isolate A5 that display two copies. Only two orthologous groups were common between the two isolates: namely, HD2 and HD1-like which are part of the putative MAT-locus in *R. irregularis* (Figure 2b). As reported in Ropars *et al.*, we observed that the two copies of the putative MAT-locus in the dikaryons were located in different contigs. One copy of HD2 and HD1-like genes were present in a long contig of the genome assembly, while the second copy was present in a much shorter contig (Figure 2c). We observed the same pattern for the other orthologous groups, where the second copy was always present in a second shorter contig (for several examples see Figure 2d).

**Figure 2.**
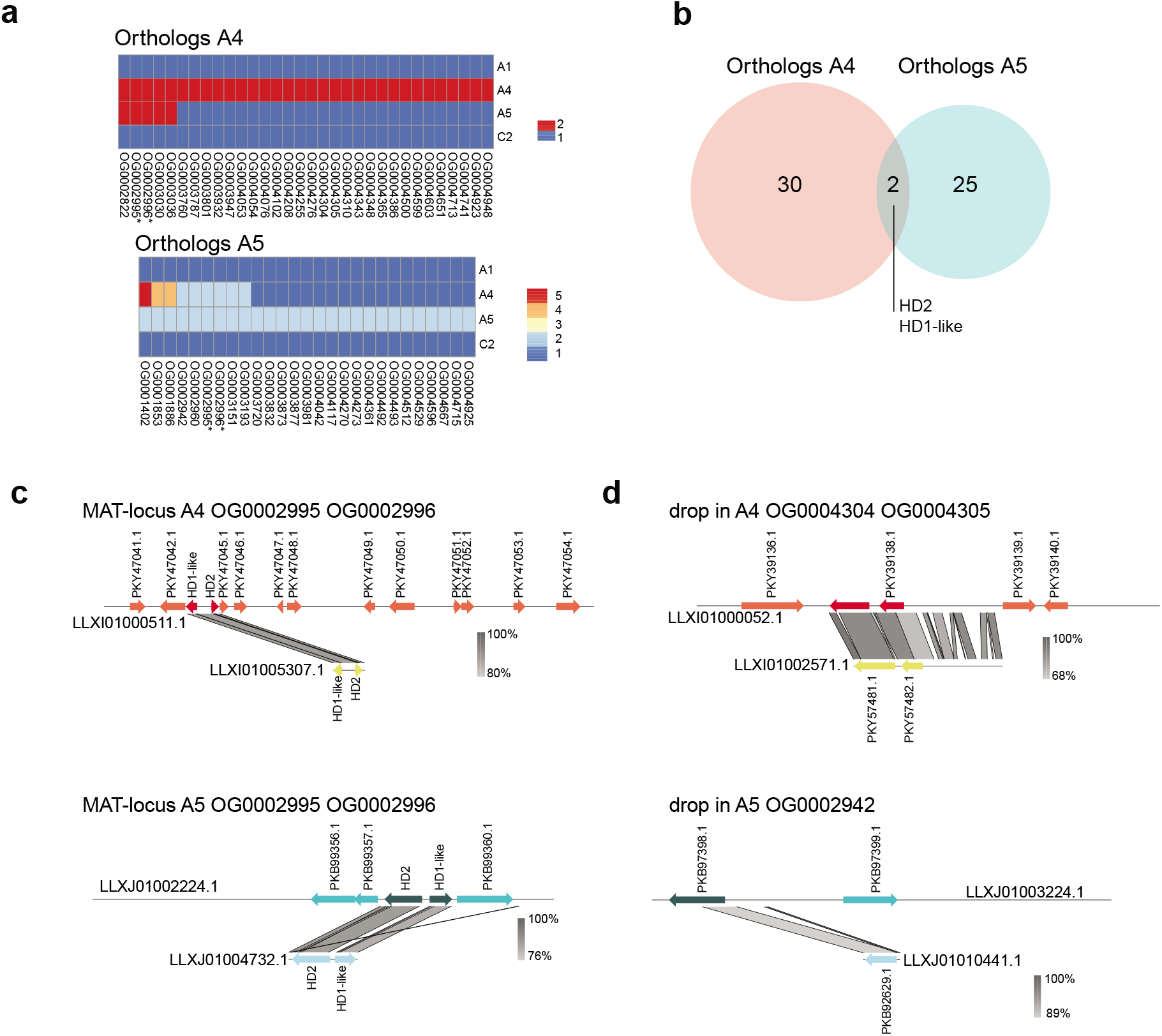
Identification of orthologs of genes present in regions showing a drop in coverage. **a,** Orthologous groups that display two or more genes in dikaryons and only a single gene in monokaryons. This analysis was performed independently on isolate A4 and isolate A5. * orthologous genes containing HD2 and HD1-like genes respectively. **b,** Venn diagram representing the number of shared orthologous groups within drop in coverage regions between isolates A4 and A5. Only two orthologous groups were shared between the isolates, they contain the putative MAT-locus genes HD2 and HD1-like. **c,** Synteny plot between the two contigs containing the different alleles of the putative MAT-locus of isolates A4 and A5. **d,** Synteny plot between the two contigs containing different alleles of other orthologous genes. Please note that the synteny figures are made from the public available annotations of each genome assembly. Differences in size of open-reading frames (ORF) among isolates are due to differences in detection of ORF on each isolate and likely could be the result of the annotation process.

To further sort paralogous genes from true orthologs, we performed a synteny analysis to compare the genomic location of the presumed orthologous genes among isolates. We identified that 12 out of 32 predicted orthologs in A4 and 16 out of 27 predicted orthologs in A5, were located in the same genomic location on the different isolates, suggesting that they should be considered as orthologs (Supplementary Table 5, for examples of inferred orthologs and paralogs see Supplementary Figure 2).

Hence, in the whole genome assemblies of dikaryon isolates, two divergent alleles were assembled into different scaffolds; one longer scaffold containing neighboring regions and a shorter scaffold without the neighboring regions. Given that the nuclei are haploid in the dikaryon isolates, two possibilities are consistent with this previous fact: The two copies could be present within the same or in different nuclei.

### Drop in coverage signatures represent nucleotype-specific haplotypes

To confirm that genes found inside drop in coverage regions are nucleotype-specific, we used sequencing reads of individual nuclei of dikaryon isolates A4 and A5 (Chen *et al.*, 2018a) to produce *de novo* single-nucleus assemblies. The *de novo* assemblies were very fragmented and incomplete (Supplmentary figure 3ad, Supplementary table 6) and their utilization was highly limited. This limitation resulted in the inability to identify some genes and some complete gene sequences. However, a reciprocal blast approach between the whole genome assembly and the single-nucleus assemblies allowed us to detect sequences in the single-nucleus assemblies corresponding to the genes detected in the whole genome assemblies.

We tested in the dikaryons if the genes identified in the drop in coverage regions were present in the form of different alleles in different single nuclei by using a reciprocal blast approach. We confirmed 9 orthologous genes to be nucleotype-specific in isolate A4 and 12 orthologous genes in isolate A5 (Supplementary Table 7). We found that the population of nuclei clustered in two groups that corresponded to the identity of the putative MAT-locus contained in each nucleus (Figure 3). This result confirms that nucleotype-specific alleles in dikaryon isolates can be identified based on genes found in drop in coverage regions and that are represented by a duplication within the genome assembly.

**Figure 3.**
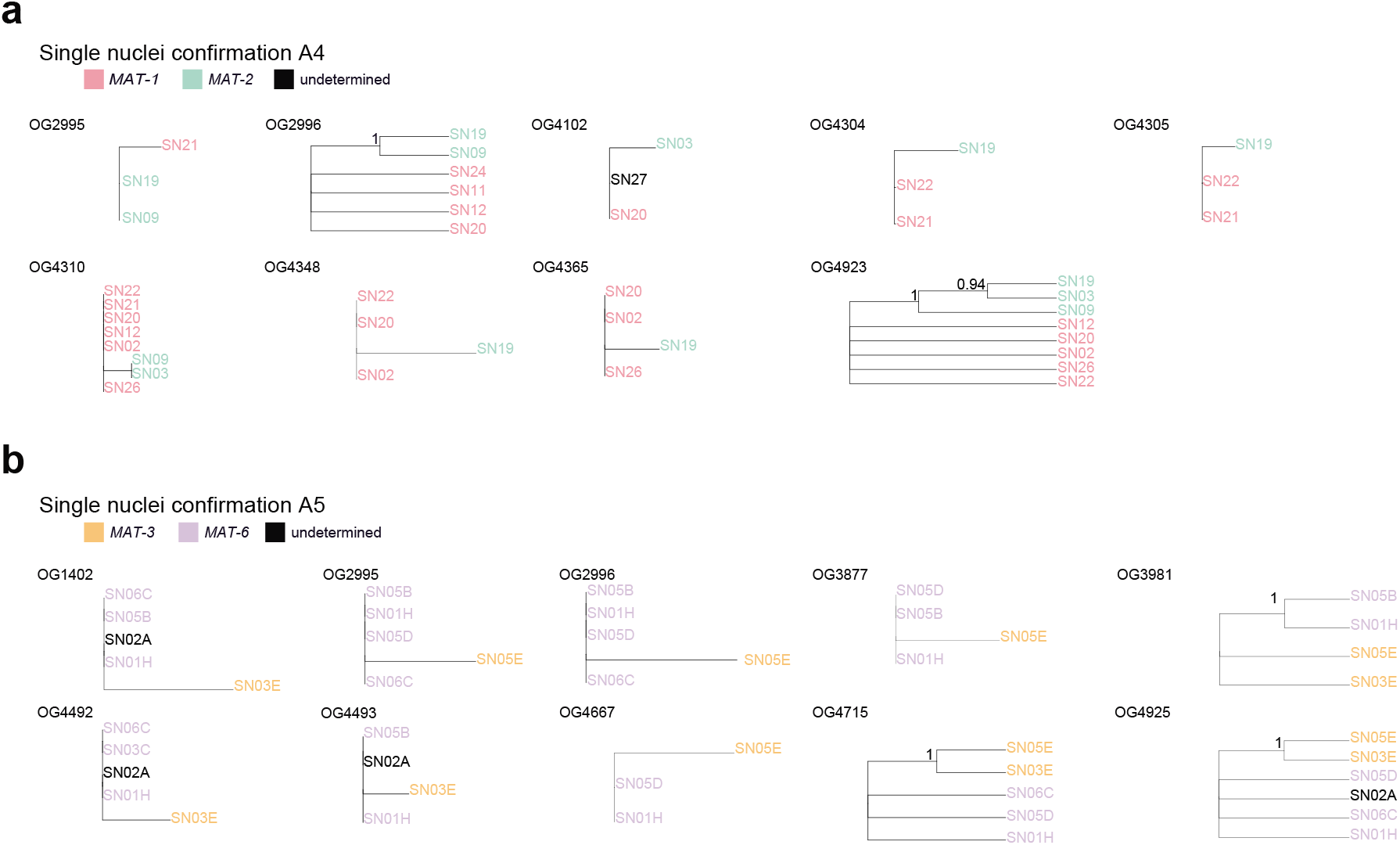
Single-nucleus sequence data confirms that genes contained in regions where a drop in coverage was observed are nucleotype-specific. Phylogenetic reconstruction of single-nucleus for genes found in regions where a drop in coverage was detected in A4 and A5 isolates. The genes are named by their membership to the orthologous groups previously defined. Branch support consisting of 100 bootstraps is shown. When only sequences from three nuclei were included, we performed an UPGMA hierarchical clustering. **a,** data for nuclei from A4 isolate. **b,** data for nuclei from A5 isolate.

### Nucleotype-specific alleles from A5 share a more recent evolutionary origin with monokaryon isolates A1 and C2 than among them

The origin of dikaryon isolates could be investigated through comparisons of monokaryon isolates that display the same alleles of the putative MAT-locus as those found in the dikaryons (Isolates A5:MAT-3/MAT-6; A1:MAT-3; C2: MAT-6). A phylogenetic reconstruction of the putative MAT-locus suggests that MAT-3 from isolates A1 and A5 are more closely related than MAT-6 from isolates C2 and A5 (Ropars *et al.*, 2016). Furthermore, genome-wide reduced genome representation phylogenetic reconstructions of several *R. irregularis* isolates indicated that isolate A5 is more closely related to isolate A1 than to isolate C2 (Wyss *et al.*, 2016; Savary *et al.*, 2018).

To confirm the previous findings, for each previously defined nucleotype-specific gene, we compared the phylogenetic relationship of the two nucleotype-specific alleles in isolate A5 isolate and in isolates A1, C2 and A4. We observed that for several nucleotype-specific genes, genes from isolate A1 clustered with one of the alleles of isolate A5, but it was not always the case in isolate C2 (Figure 4a).

**Figure 4.**
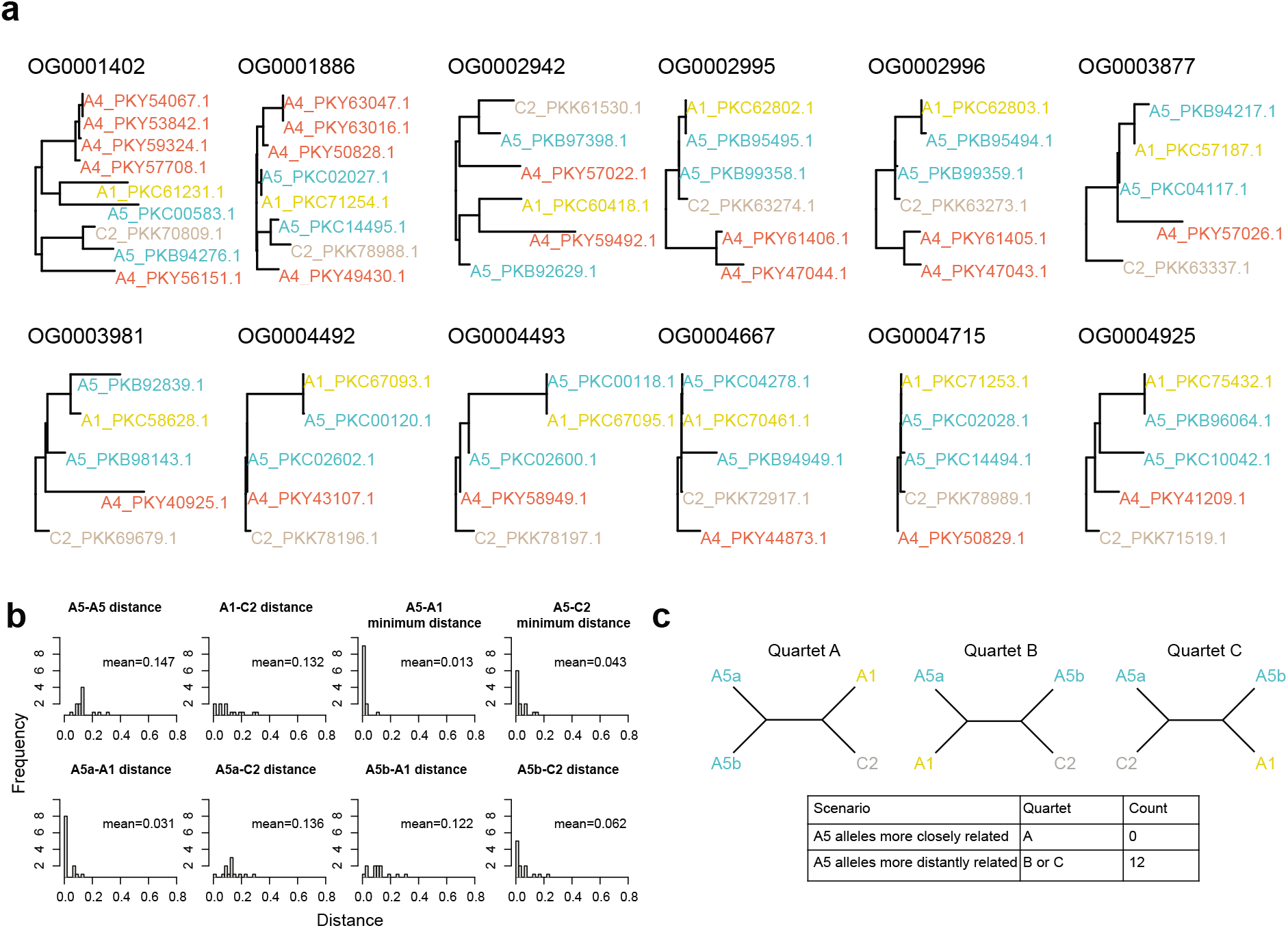
Nucleotypes from isolate A5 share a more recent evolutionary origin to isolates A1 and C2 than among them. **a,** Phylogenetic reconstruction of nucleotype-specific alleles in isolate A5 and its orthologs in isolates A1, C2 and A4. **c,** Average genetic distances between the different nucleotype-specific alleles from two alleles from isolates A5 and their homologs in isolates A1 and C2. We show histograms representing the genetic distance between the two nucleotype-specific alleles of isolate A5 and their homologues in isolates A1 and C2. For comparisons between A5 and C1 or A1 we used the minimum distance of the two alleles from A5. **d**, Scenarios of genetic similarity between the two A5 alleles and alleles from A1 and C2.

We then analyzed the genetic distance of each of the 12 nucleotype-specific genes independently between the two alleles of isolate A5 and the homologous allele in isolates A1 and C2. The mean nucleotide distance between the two A5 alleles was 0.147. The mean distance between A1 and C2 was 0.132. In contrast, the mean of the minimum distance between an allele of isolate A5 and isolate A1 was 0.013 and between A5 and C2 was 0.043 (Figure 4b). We did not observe any case where the two A5 alleles clustered together, instead we observed that for all 12 nucleotype-specific gene the two A5 alleles were more similar to the allele from isolate A1 or C2 (Figure 4c). The mean distances calculated between alleles in this study are much higher than average distances calculated on the whole genome between different isolates (Chen *et al.*, 2018b), reflecting our selection criteria for nucleotype-specific regions. As each of the A5 alleles was closer to A1 or C2, instead of the two A5 alleles being most similar, this indicates that the alleles of A5 share a more recent evolutionary origin with these monokaryons than the two alleles within A5.

### Identification of reciprocal recombination between nucleotype-specific haplotypes in isolate A5 using single-nucleus data

Knowledge about nucleotype-specific haplotypes of dikaryon isolate A5 and their orthologs in isolates A1 and C2 allowed us to test whether parasexual or sexual genomic signatures could be identified in dikaryon isolate A5 (Figure 5a). We scanned the different nucleotype-specific-haplotypes for the detection of recombination events within the two haplotypes of isolate A5. Comparison of haplotypes issued of the short-read whole genome assemblies from isolates A1, C2 and the two haplotypes of A5 showed that each nucleotype-specific sequence from isolate A5 was highly similar to either isolate A1 or C2, but we did not identify any recombination events within the haplotypes (Supplementary Figure 3).

**Figure 5.**
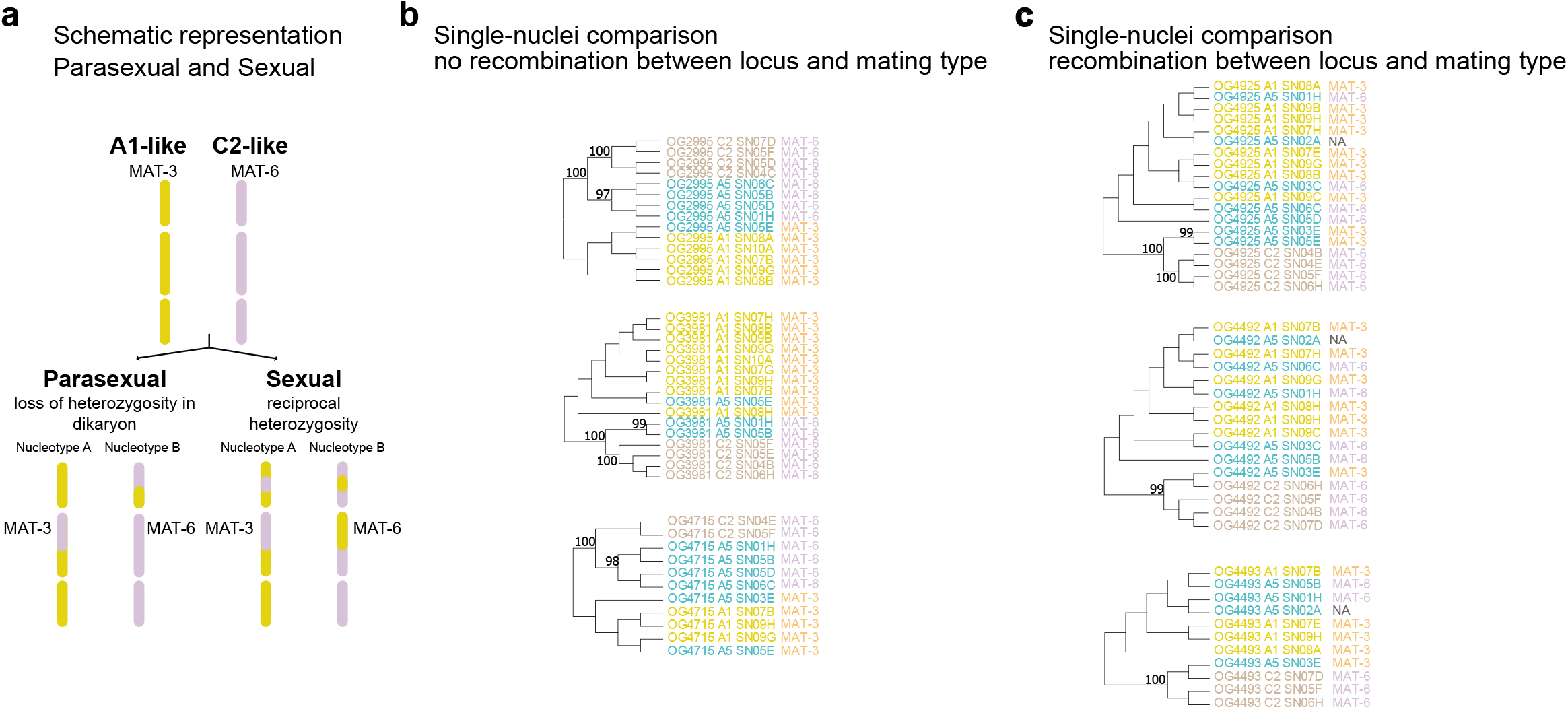
Recombination events in nucleotype-specific haplotypes in A5 isolate. **a,** Schematic representation of possible outcomes after fusion of two different isolates. Please note the schema illustrates different contigs, separated by blank lines and no different chromosomes. **b,**Phylogenetic relationship of different nucleotype-specific alleles among different nuclei from A1, A5 and C2 isolates. Cases where no recombination was detected. Nuclei having the same MAT-type clustered together. **c**, Phylogenetic relationship of different nucleotype-specific alleles among different nuclei from A1, A5 and C2 isolates. Cases where recombination was detected. Nuclei having the same MAT-type did not clustered together. We performed 100 bootstraps for the branch support of all phylogenetic constructions.

To further assess the potential for clonal relationships between the two nucleotypes within A5 and isolates A1 and C2, we compared the nucleotype-specific haplotypes on the single-nucleus assemblies from isolate A5 and their orthologs on the single-nucleus assemblies of isolates A1 and C2 (Supplementary Table 8). The difference with the previous analysis is that with the single-nucleus data, we are able to identify the identity of the putative MAT-type of the A5 haplotypes. We found that for several nucleotype-specific genes (i.e. OG2995, OG3981 and OG4715), the sequences from isolates A5 (MAT-3 type) and A1 (MAT-3 type) clustered together (Supplementary Figure 4, Figure 5b). However, we found that for other nucleotype-specific genes (OG4925, OG4492 and OG4493), the sequences from isolate A5 (MAT-3 type) clustered with the sequences from isolate C2 which has a MAT-6 type (Supplementary figure 5, Figure 5c). The alignments on these nucleotype-specific genes show that A5 nuclei with MAT-3 have similar, but not identical alleles as C2 nuclei (MAT-6 type). In the same way, A5 nuclei with MAT-6 type harbor alleles similar to those of A1 nuclei (MAT-3 type). These results demonstrate that the A5 isolate harbors nucleotypes with regions highly similar to isolates A1 and C2, but that the A1-like alleles are not always found in the same nucleus. The presence of reciprocal recombinant nucleotypes in isolate A5, involving isolates sharing the same MAT-type, strongly suggest that isolate A5 results from a sexual and not parasexual event between isolates similar to A1 and C2.

### Confirmation of reciprocal recombination events in isolate A5 using long-read genome assemblies

We compared single-copy orthologous genes among the A1, C2 assemblies and the phased assembly of isolate A5, which is divided in a primary assembly and an haplotig assembly.

The long-read assemblies were more complete and contain longer scaffolds than the short reads assemblies (Supplementary Figure 3bcef). The A5 phased assembly consisted of a primary assembly of 392 scaffolds and an haplotig assembly on 292 of the primary scaffolds. The primary assembly was 115Mb long and the haplotig assembly was 19.2Mb (Chaturvedi *et al.*, 2021).

We identified 1250 single-copy orthologs among isolates A1, C2, the A5 primary assembly and the A5 haplotig assembly (Supplementary Table 9). We then focused only in the orthologs were one of the genes of A5 clustered with either A1 or C2. The majority of the genes sequences were very similar among all the isolates, however we identified 44 orthologous groups where one of the haplotype genes of A5 cluster with either isolate A1 or C2 (23 orthogroups A1-A5haplotig and 21 orthogroups A1-A5primary). We observed that for several contiguous regions, an orthogroup displayed the A5primary gene clustering with A1 and in another orthogroup, contained in the same contiguous region, the A5 primary gene clustered with the C2 gene (Figure 6), confirming that reciprocal recombination signatures are found within continuous haplotypes (Supplementary Table 9).

**Figure 6.**
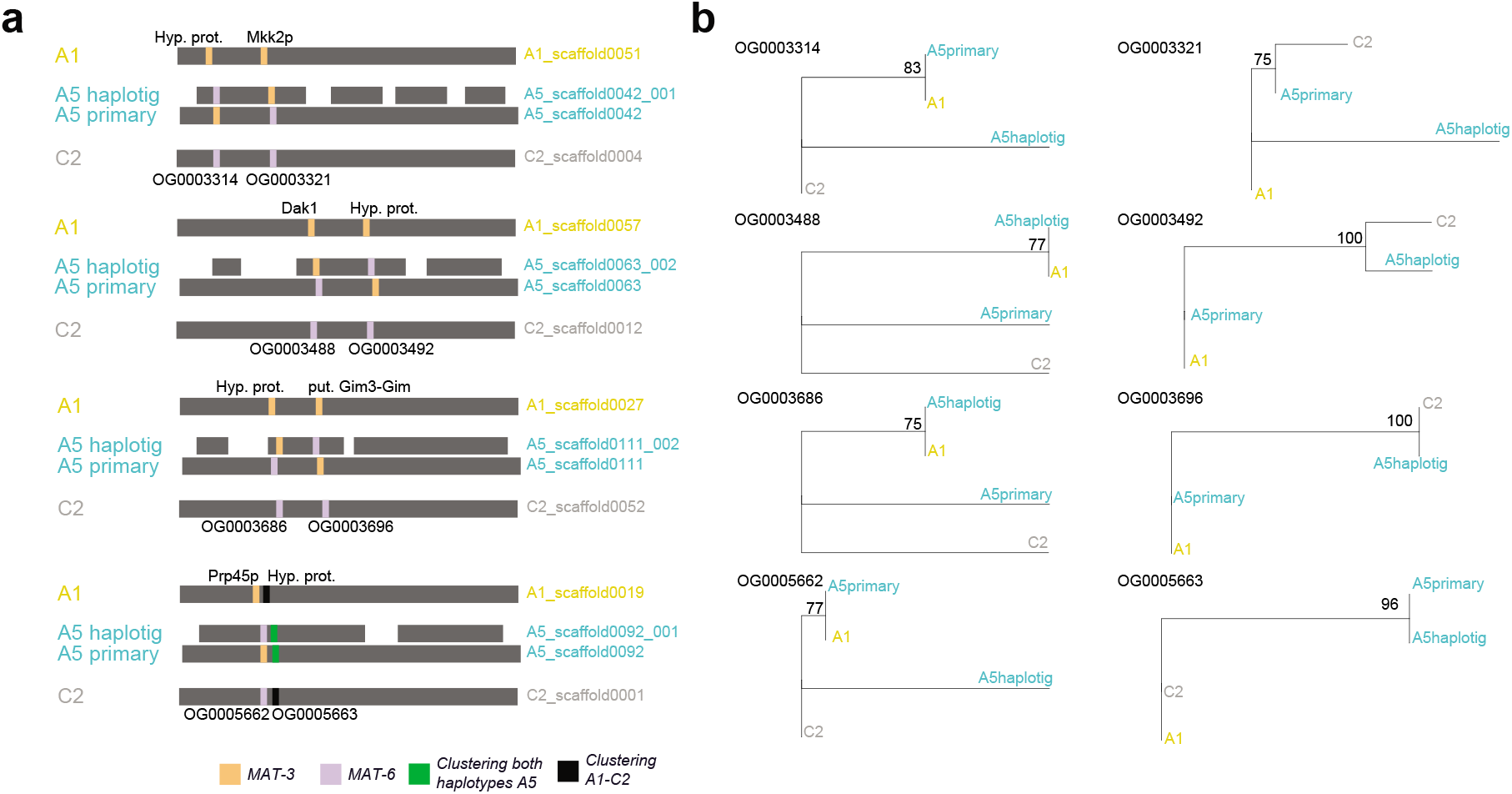
Recombination events identified in continuous haplotypes issued from the long-read genome assembly of dikaryon isolate A5. **a** Schematic representation of recombination events within continuous haplotypes. We highlight single-copy orthologous genes represented by different colors depending if they cluster with a gene of isolate A1 MAT-3 (yellow) or C2 MAT-6 (violet). Genetic recombination is demonstrated when within a continuous haplotype (grey background) a change in clustering pattern between an A5 haplotype and isolate A1 or C2 is observed. We can also observe that for few cases the two haplotypes of A5 clustered together (green). **b** Phylogenetic reconstructions of the different single-copy orthologous groups identified within the same haplotype. We can observe that the clustering pattern change within the same haplotype (i.e. A1-A5primary and C2-A5primary). We performed 100 bootstraps for the branch support of all phylogenetic constructions.

## Discussion

In this study, we used short and long read published sequencing data to demonstrate that signatures of reciprocal recombination can be detected in genomes of dikaryon AMF isolates that display different putative MAT-types. The signatures of reciprocal recombination are compatible with a sexual reproduction life cycle, rather than a parasexual reproduction mechanism. These results suggest that after all, AMF are not that scandalous.

In the absence of experimental evidence that isolates with different putative MAT-types are involved in sexual reproduction, genomic signatures of recombination can help us to understand whether sexual or parasexual reproduction are involved in the AMF life cycle. On one hand, parasexual reproduction genomic signatures involves gene-conversion, which result is the loss of heterozygosity (Forche *et al.*, 2008). On the other hand, reciprocal recombination associated with a putative sex-determining region (MAT-type) is a genomic signature of sexual reproduction, although gene-conversion could also happen in sexual reproduction (Sun *et al.*, 2012). It could be plausible that parasexual recombination could lead to a reciprocal recombination pattern by fusion of two monokaryons, mitotic reciprocal recombination, followed by reciprocal chromosome loss. However, the presence of a complete meiotic machinery, the presence of compatible putative MAT-types, and the upregulation of genes involved in a putative mating response when two isolates co-exist (Mateus *et al.*, 2020, 2021), suggest that a plausible hypothesis explaining the results observed in this study is a reciprocal recombination pattern detected in dikaryons, the result of sexual reproduction.

We identified few regions that display recombination patterns within continuous haplotypes in the dikaryon isolate A5. The principal reason, is that we did not evaluate all the different genes present, but only the genes that display only 1 copy in an haploid phased contig. We only compared single-copy orthologs among the different assemblies, to avoid the confounding effect of paralogy on our results. In consequence, the examples of reciprocal recombination shown in this study, are only a small subset of the total plausible recombination spots present in their genomes. With the information of nucleotype-specific haplotypes identified with the drop in coverage analysis, we found that nucleotypes of isolate A5 were as little 1% diverged from isolate A1 and 4% diverged from isolate C2, suggesting that isolates sharing the same MAT-type as the monokaryon isolates A1 and C2 are closely related ancestors from which the dikaryon A5 arose. Interestingly, with the long-read data, we identified several loci where the A1 and C2 genes were more closely related among them than to the alleles of isolate A5. This suggest, that there could be other evolutionary forces shaping AMF genome evolution. Transposable elements (TE) have been reported to be responsible of genome duplications, inversions, insertions and deletions (Daboussi & Capy, 2003). In nature, AMF co-exist with different conspecifics and they co-exists aswell with different host-plants. We could speculate, that gene trajectories within AMF populations could be also driven by TE-mediated horizontal gene transfer events. However, an important sampling of AMF genomes is necessary to better understand the genome architecture features and life-history of the different gene families (Badet & Croll, 2020).

An intriguing question, is how only two coexisting nucleotypes and a well-orchestrated mechanism such as meiosis could be achieved without the presence of a single-cell stage in the AMF life cycle. A genetic bottleneck, where only 1 or 2 nuclei co-occur has never been detected in AMF. Despite, the last feature, two studies that evaluated nuclear imbalance in dikaryon isolates show that nuclei ratio among single-spore lines are conserved despite multiple culturing for years (Robbins *et al.*, 2021) or are stable when exposed to different host plants (Kokkoris *et al.*, 2021). These studies suggest that a nuclei regulation mechanism should exist and could be responsible of the stability of the nucleotypes in the dikaryons.

In this study, we compared haplotypes within the dikaryon A5 to monokaryon isolates (A1 and C2) that contain the same putative MAT-types as the dikaryon A5 to identify the recombination patterns.

We were able to analyze these signatures on the A5 dikaryon, but not on the dikaryon A4, because there are no available genomic assemblies of monokaryon isolates displaying the same putative MAT-type as in Isolate A4. We hypothesized that MAT-type compatibility exists between the two different putative MAT-types identified in the dikaryon isolates. However, to date there is no direct experimental evidence that different putative MAT-types could be compatible. An indirect evidence of compatibility between different putative MAT-types was found, when two different isolates harboring different putative MAT-types elicited a putative fungal mating response (Mateus *et al.*, 2020, 2021). However, Mateus et al., did not test if the mating of two co-existing isolates did took place (Mateus *et al.*, 2020). In consequence, in order to experimentally identify mating and MAT-type compatibility, crossing experiments between strains harboring different putative MAT-types, including transcriptome and recombinant progeny analyses should be performed.

Our drop in coverage approach allowed us to identify divergent nucleotype-specific genes in dikaryon isolates situated in different contigs of the short-read whole genome assemblies. This approach differs from previous approaches of global intra-isolate divergence assessment that measured the number of SNPs (Chen *et al.*, 2020) or poly-allelic sites (Wyss *et al.*, 2016). Although the comparison of both types of measurements gave similar information (intra-isolate divergence), their comparison should be carefully addressed as their methodology and the types of sequences compared are different. While SNPs are best identified in low divergence regions, where reads can be confidently mapped to the same contig, the sequences with large genetic differences, detected with the drop in coverage approach, are highly divergent to the point that they are assembled in different contigs in the same genome assembly. Consequently, genetic divergence should be higher in the regions with large genetic divergence than when the two alleles are collapsed in the genome assembly, resulting in sequences displaying several SNPs. Contrary, to the drop in coverage analysis using short-read data, the phased haplotypes from long-read sequencing, display genes that differ in several SNP’s between the two dikaryon haplotypes.

Inter-nucleus recombination has been previously reported (Chen *et al.*, 2018a), although the robustness of the analysis has been questioned. The application of strict filtering parameters such as removal of heterozygous sites in haploid nuclei, duplicated regions of the genome, and low-coverage depths base calls results in an extreme loss of the signal of recombination (Auxier & Bazzicalupo, 2019). Although some of these limitations, as coverage depth and filtering-out heterozygous sites were addressed (Chen *et al.*, 2020), other limitations such as replicability (recombination events shown in several nuclei sharing the same putative MAT-type) and issues related to the whole-genome amplification step as the formation of chimeric sequences (Yilmaz & Singh, 2012), allelic drop-out (Lauri *et al.*, 2013) and SNP miscalling (Ning *et al.*, 2014) are inherent limitations of the analysis of any single nucleus amplification data. Furthermore, the reported evidence of large inter-nuclei recombination (Chen *et al.*, 2018a, 2020) does not fit the observations about a monokaryon – dikaryons organization. In Chen *et al.*, 2018a, the selected examples of genotypes presented in figure 3 show 4 nuclei per isolate, which represent four different nuclei for SL1, A4 and three different nuclei on A5. The same pattern can be observed in Supplementary Figure 5 from Chen *et al.*, 2020, the examples of genotypes show 7 out of 8 different nuclei in isolate A5, at least 7 different nuclei in isolate A4 and at least 5 different nuclei in SL1. The presence of repeated inter-nuclear recombination in dikaryons without prior crossing with other isolates, as observed in the single-nucleus genotypes shown in Chen et al. (2018a, 2020) would result in heterokaryons with more than two types of nuclei. But this does not appear to be the case for *R. irregularis* (Ropars *et al.*, 2016; Chen *et al.*, 2018a; Masclaux *et al.*, 2019; Auxier & Bazzicalupo, 2019).

Maintaining monokaryon and dikaryon isolates within the same natural population suggests that both forms are stable over time. Rather than a promiscuous mixing between isolates via anastomosis, a mechanism of recognition that involves the putative MAT-locus seems to regulate which isolates can form a dikaryon (Corradi & Brachmann, 2017). The fact that nucleotype-specific haplotypes from isolate A5 are more closely related to isolates A1 and C2, and that A5 nucleotypes display recombination, suggests that isolates sharing the same putative MAT-type as A1 and C2 could be the origin of a recombining dikaryon isolate. However, we cannot discard that these findings could apply to another step of the AMF life cycle. It could also be possible that a stable A5 isolate could segregate producing recombined monokaryons that share the same putative MAT-type as isolates A1 and C2, that can disperse and then fuse again to form stable dikaryons and complete a life cycle which involves recombination. It then becomes crucial to experimentally confirm if monokaryon isolates having different putative MAT-types could generate a dikaryon-like form and if a dikaryon isolate could segregate into recombining monokaryon isolates.

Understanding the life cycle of AMF could have an enormous impact in the generation of AMF genetic variability. The generation of diverse AMF monokaryons or dikaryons could be used to generate variants that enhance plant growth and have an enormous potential in agriculture (Sanders, 2010).

## Acknowledgments

We would like to thank Daniel Croll for his feedback on a preliminary version of this manuscript. We specially thank Bart Nieuwenhuis and his group for the constructive feedback made on the Biorvix version of this manuscript. This study was funded by the Swiss National Science Foundation (Grant number: 31003A_162549 to IRS).

## Author contributions

IDM designed and supervised all the analyses. MMSN developed the drop in coverage detection algorithm. JC identified the genes present in the regions displaying drop in coverage. BA developed the genetic relatedness analysis on the nucleotype-specific haplotypes. SL provided valuable comments during all the process. IDM, performed all the bioinformatic analyses, identified the nucleotype-specific genes, performed the phylogenetic analysis, gene-retrieval from genome assemblies and the recombination detection. IDM and IRS wrote the manuscript. All the authors provided valuable contributions and modifications of the manuscript.

## Data availability

All data analyzed in this manuscript its available in the NCBI repository and the respective accession identifiers could be found in Supplementary table 1. The custom code developed for the identification of drop in coverage regions could be found in Supplementary file 1.

