## Supplementary material for "Reciprocal recombination reflects sexual reproduction in symbiotic arbuscular mycorrhizal fungi"

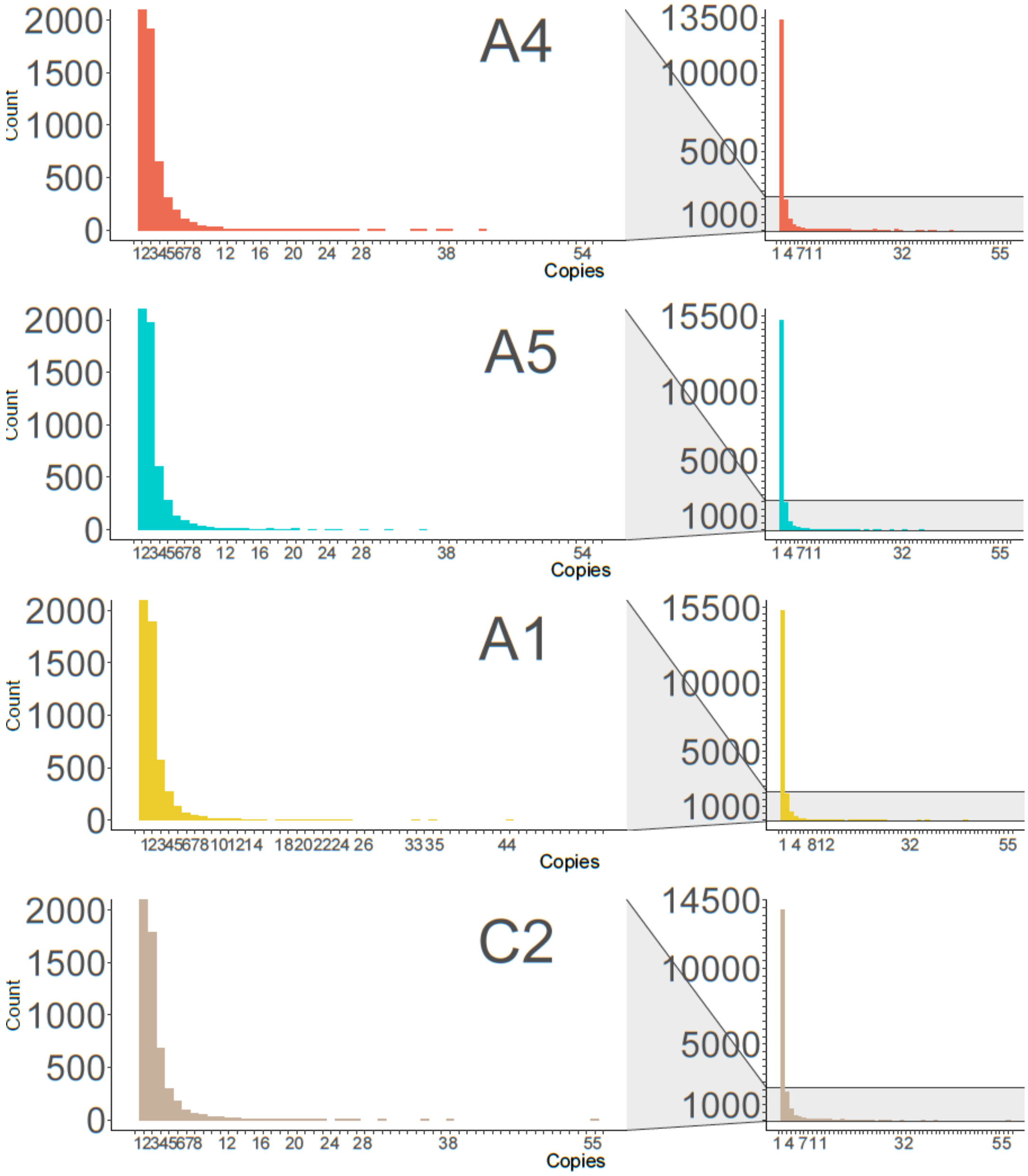


**Supplementary Figure 1. Number of copies of each orthologous group within each isolate.** Number of copies of different orthologous groups found within the genome of each isolate.

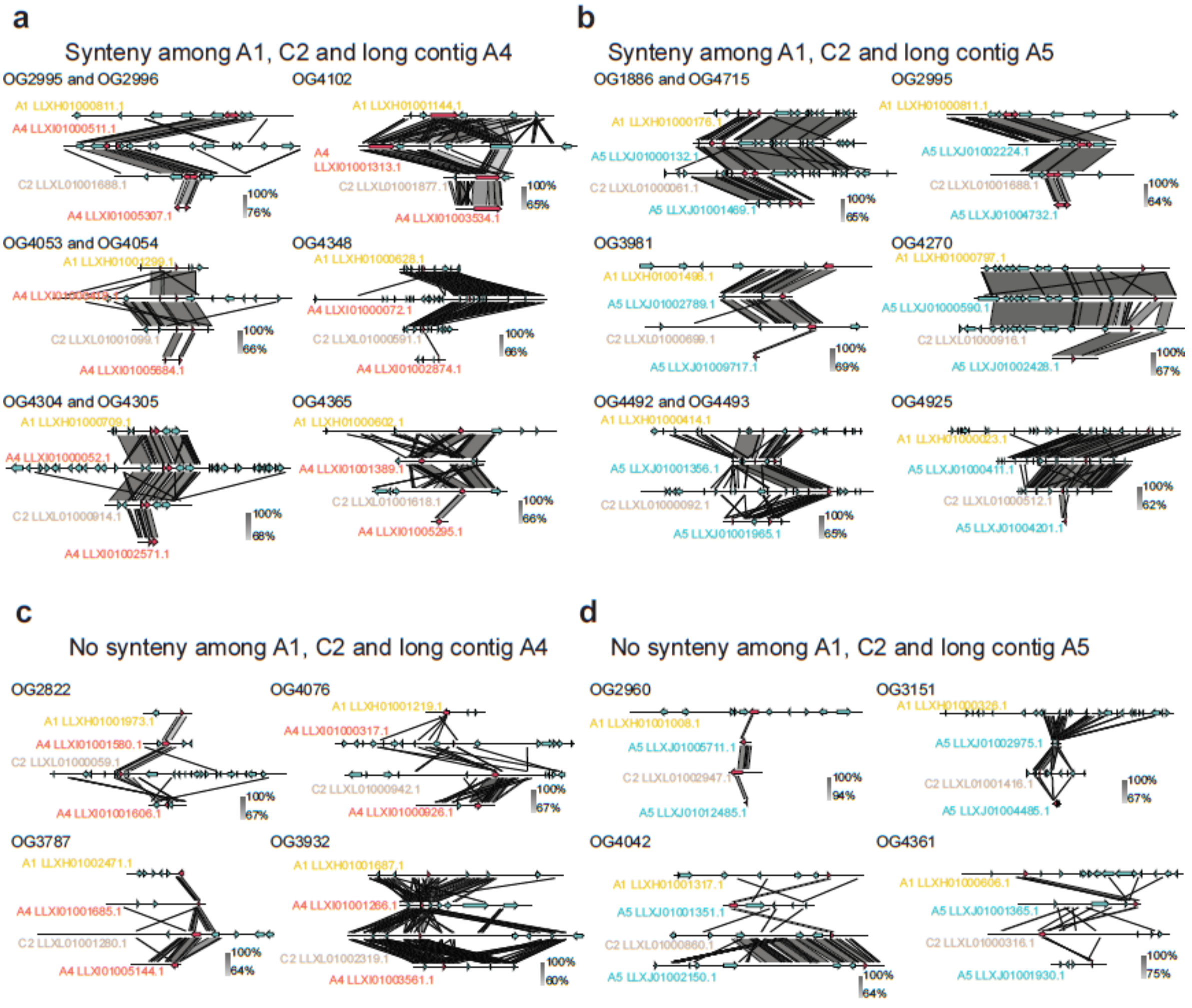


**Supplementary Figure 2. Synteny among candidate nucleotype specific regions in isolates A1, A4, A5 and C2. a,b** We show examples of orthologous genes on different isolates that are situated in the same genomic location in isolate A4 and isolate A5 respectively. This is evidenced by the blast homology (Grey zones linking the different contigs) shown in surrounding regions of the focal gene (in red). **c,d** Examples of paralogous genes on different isolates where there genomic location of the gene is not the same. Evidenced by the lack of homology (grey zones linking the different contigs) of the surrounding regions of the focal gene (in red).

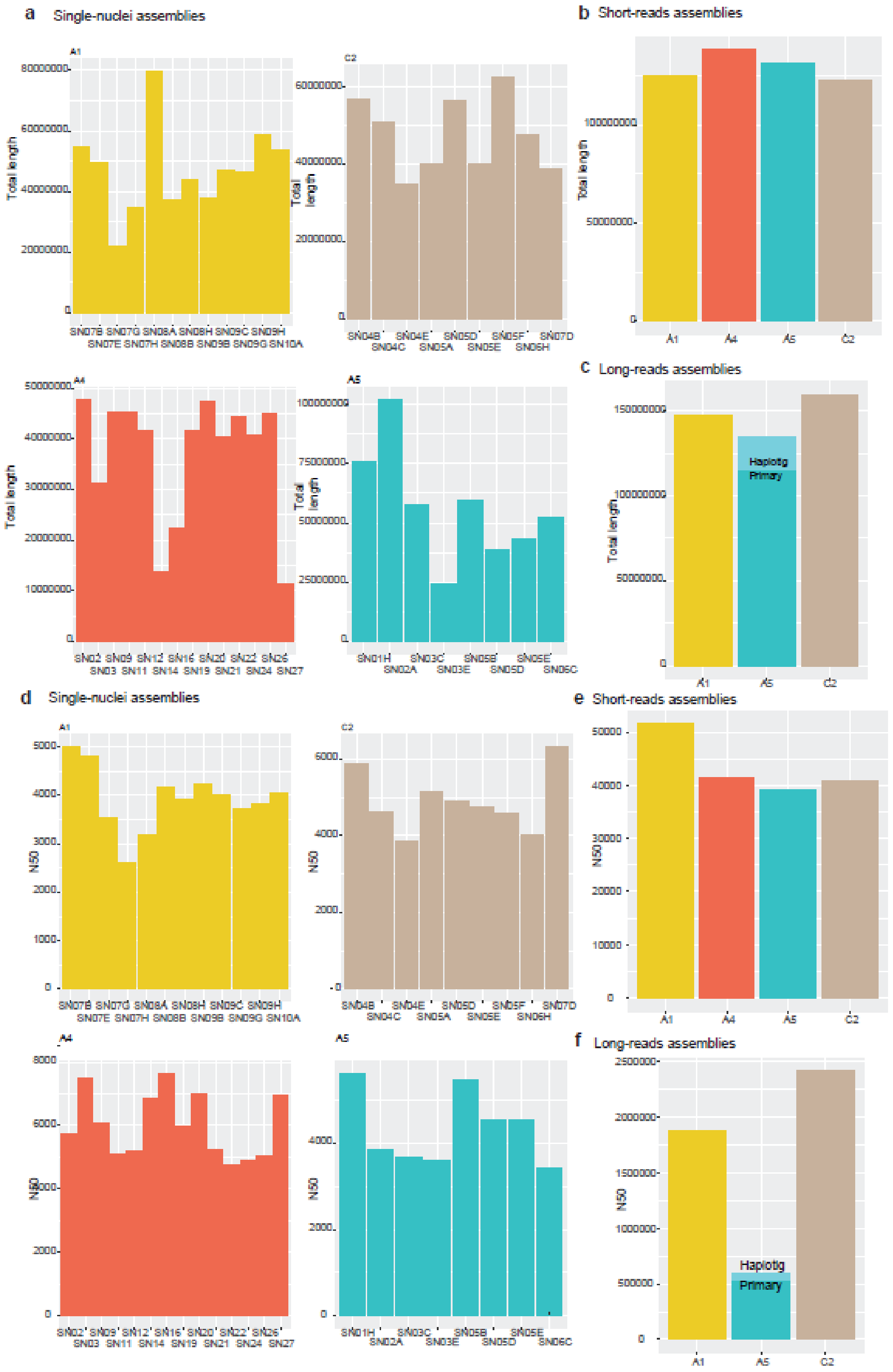


**Supplementary Figure 3. Total length and N50 statistics of the genome assemblies used in this study. a,d** Single-nuclei assemblies, **b,e** short-reads assemblies, **c,f** long-reads genome assemblies.

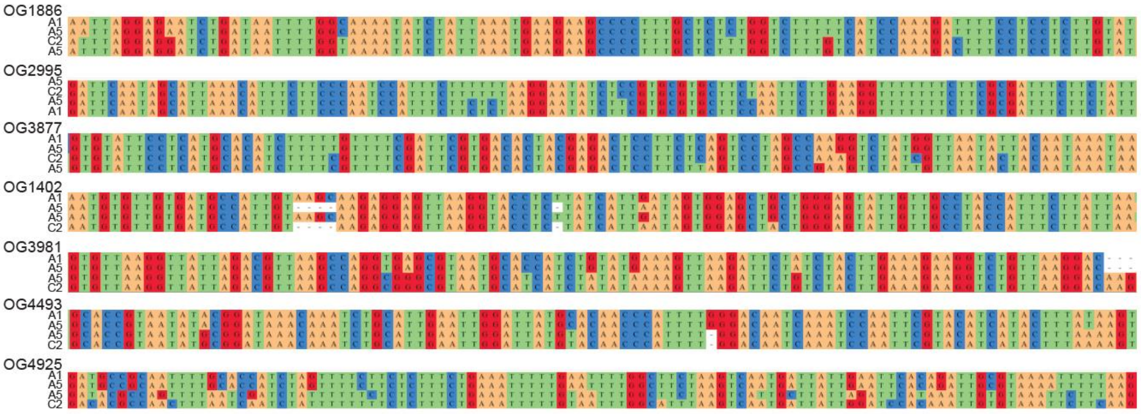


**Supplementary Figure 4. Multiple sequence alignment of nucleotype-specific orthogroups of isolate A5 and isolates A1 and C2.** Sequences are issued from the short-reads genome assemblies. The sequences shown are collapsed and do not represent the total length of the genes.

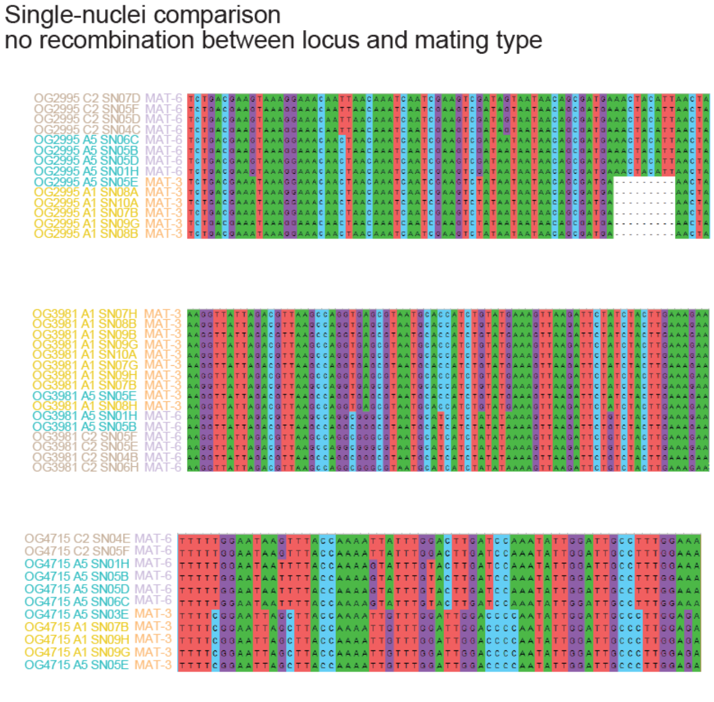


**Supplementary Figure 5. Multiple sequence alignment of nucleotype-specific orthogroups of single-nuclei of isolate A5 and isolates A1 and C2.** This example represents the case when no recombination is identified. Please note that MAT-3 or MAT-6 sequences cluster together. The sequences are issued from the single-nuclei genome assemblies. The sequences shown are collapsed and do not represent the total length of the genes.

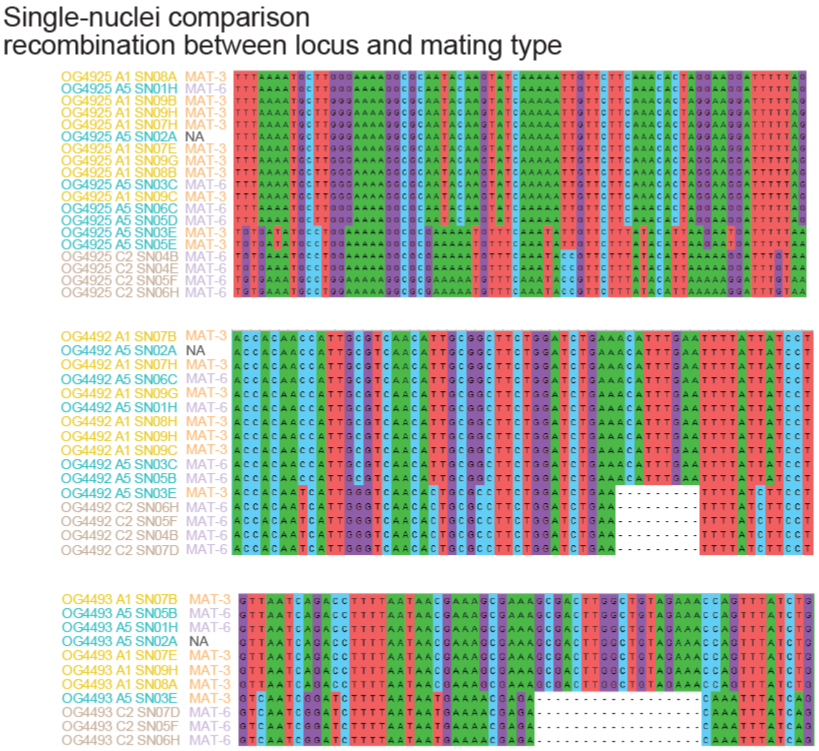


**Supplementary Figure 6. Multiple sequence alignment of nucleotype-specific orthogroups of single-nuclei of isolate A5 and isolates A1 and C2.** This example represents the case when recombination is identified. Please note that MAT-3 or MAT-6 sequences do not cluster together. The sequences are issued from the single-nuclei genome assemblies. The sequences shown are collapsed and do not represent the total length of the genes.
